# Acute and sub-chronic toxicity of aqueous stem-bark extract of *Annickia polycarpa*

**DOI:** 10.1101/2023.09.14.557718

**Authors:** Nathaniel Lartey Lartey, Eric Nana Yaw Nyarko, Henry Asare-Anane, Stephen Antwi, Laud Kenneth Okine

## Abstract

**Background and aim:** The lack of standardization and scientific validation of the use of most plant extracts leads to toxicity problems. The stem bark extract of *Annickia polycarpa* has been shown to possess antioxidant, antidiabetic, analgesic, and anti-colitis effects. However, nothing is known about the effects of the prolonged use of the extract. Thus, we investigated the acute and sub-chronic toxicity of the aqueous stem bark extract of (APE) in Sprague-Dawley rats.

**Experimental procedure:** The LD50 and sub-chronic toxicity of APE (20 mg/kg, 100 mg/kg, and 500 mg/kg) was studied over 3 months in Sprague-Dawley rats. Serum alkaline phosphatase (ALP), alanine transaminase (ALT), direct bilirubin and creatinine were measured after 3 months of treatment. Hematological analysis and urinalysis were also performed. Also, the effect of APE on blood clotting time and pentobarbital-induced sleeping time were determined at the termination of treatment. Finally, histological analysis was done on liver, kidney, lung, and heart after hematoxylin-eosin staining of tissue cross-sections at the termination of treatment.

**Results and conclusion:** The LD50 of APE in rats was higher than 5000 mg/kg with no observable signs of toxicity. APE also showed no hematotoxic effect when used consistently for 3 months. Additionally, APE had no adverse effect on the liver and kidney evidenced by a lack of effect on serum biochemical parameters (ALP, ALT, direct bilirubin, and creatinine), urinalysis as well as the tissue morphology of these organs. No adverse morphological effects were also observed with the heart muscle cells. However, APE showed mild selective toxicity to the lung characterized by alveolar space closing and interstitial fibrosis and alveolar septa thickening. These results indicate that APE is generally non-toxic at the tested doses but shows mild selective pneumotoxicity when used over a long period of time.

## Background

The use of natural products and herbal preparations for the treatment of diseases has been with man for a long time, but it’s only until some decades ago when studies began to be conducted on them ^1^. An estimated 60 to 95% of rural Africans are said to be reliant on traditional herbal preparations for their primary health care needs ^2,3^. Therefore, in developing countries with meagre resources, herbal preparations have an ever-important role to play in preventing the complications of lifelong diseases such as diabetes mellitus ^4^.

The high cost of treatment and management of many disease conditions with orthodox medications coupled with undesirable side effects has caused people, particularly rural folks, to resort to the use of medicinal plants preparations because they are cheaper and easily accessible ^5^. One major problem with the use of herbal preparations is the lack scientific validation of the safety, efficacy and standardization of these products, which may result in possible overdosing and consequent toxicity, thereby putting consumers at risk. The stem bark extract of *Cassia sieberiana* used in North-eastern Nigeria for the treatment of jaundice, for example, has been proven to be hepatotoxic at lower doses of 20-60 mg/kg and nephrotoxic at a higher dose of 180 mg/kg ^6^. This observation is at variance with the famous assertion that plant extracts are non-toxic and thus the need to conduct toxicity studies on them.

*Annickia polycarpa* is a small to medium-sized tree also known African yellow wood that grows up to 20 m tall ^7^. The stem of *Annickia polycarpa* is typically straight and cylindrical and up to 40 cm in diameter, with no buttresses ^7^. Its bark is usually smooth to slightly rough or fissured. The stem bark has been used to treat fever, including malaria ^7,8^. Also, the stem bark extract of *Annickia polycarpa* (APE) possesses antioxidant activity mainly by scavenging free radicals ^9^. In addition, its antibacterial, antifungal and antitrypanosomal effects have been reported ^8,10–12^. Palmatine, a potent anti-inflammatory agent, has been isolated and characterized from the stem bark ^13^.

Discovery of traditional medicines which are relatively efficacious, display a rapid onset of drug action, produce little or no side effects, readily available and cheaper may prove beneficial in the search for better, affordable, and more effective ways of managing diseases. There are anecdotal claims of the use of *A. polycarpa* for the management of diabetes mellitus in Ghana, which has been validated scientifically in our laboratories ^9^. We additionally showed that APE possesses anti-inflammatory, antioxidant and anti-colitis effects ^14^. However, there is the paucity of available scientific data as to its use as a safe herb to regulate its rational use. The study was conducted to assess the acute and sub-chronic toxicity of aqueous stem bark extract of *A. polycarpa* in laboratory animals. We show here that APE is generally non-toxic except for mild selective toxicity to the lung when used repeatedly for a long time.

## Materials and Methods

### Materials

#### Reagents and Chemicals

Alanine aminotransferase (ALT), alkaline phosphatase (ALP), creatinine, direct bilirubin and glucose assay kits were obtained from ELI Tech (Puteaux, France). Urine test strips (UroColor^TM^10) were supplied by Standard Diagnostics Inc. (Kyonggi-do, S. Korea). All other chemicals and reagents used were obtained from FLUKA (Zofingen, Switzerland).

#### Plant Material

The stem bark of *Annickia polycarpa* was collected from Begoro in the Eastern Region of Ghana (<u>6°23′N 0°23′W</u>) and authentication was done by the Plant Production Department (PDD) of the Centre for Plant Medicine Research (CPMR), Ghana. It was subsequently deposited with the herbarium of the PDD and given voucher specimen number of 01/2015.

#### Experimental Animals

Sprague-Dawley rats (180-200 g) were obtained from the Animal Experimentation Unit, CPMR. All animals used in this study were handled following the Guide for the Care and Use of Laboratory Animals. Stainless steel metal cages with wood shavings as bedding were used to house the animals. Standard animal feed obtained from GAFCO (Tema, Ghana) was provided for the animals and allowed free access to sterilized distilled water. The environment of the experimental animals was characterized by a temperature of (22 ± 2 °C) temperature and humidity (70 ± 4%) under alternating 12-hour periods each of light and darkness. Maximum of 6 animals were kept in a cage sized (20.3 cm width × 28.7 cm length × 17.3 cm height) thus ensures enough space for free movement. The Ethical and Protocol Review Committee of the College of Health Sciences, University of Ghana, provided ethical clearance for this work with Protocol Identification Number: CHS-Et/M.4 – P 3.10/2015-2016.

### Methods

#### Preparation of Plant Material

The plant material was prepared as has been described previously ^9,13,14^. Briefly, the fresh stem bark of *A. polycarpa* (AP) was subjected to air-drying at a temperature of 25^°^C for 1 week. The dried bark was milled into a powdered form, weighed, and boiled using water (20% w/v) for thirty (30) minutes. The extract was filtered through cotton wool after cooling and the filtrate concentrated using a rotary evaporator and lyophilized into powder by a freeze dryer (EYELA, Tokyo Rikakikai Co. Ltd, Japan). The weight of the freeze-dried extract of AP, now referred to as APE, was determined and kept in a desiccator at 4-8 ^°^C.

#### Acute toxicity study

A single oral dose at 5000 mg/kg (2 ml/rat) of APE was administered to six rats. Physical signs of toxicity and death of the rats were observed for 24 hours. Animals that survived were observed for an additional 13 days for symptoms toxicity such as piloerection, and defects in lachrymatory, locomotory and respiratory activities.

#### Sub-chronic toxicity study

The animals were divided into four groups of six rats each and placed into four separate metal cages. Animals belonging to group one served as the control group and were given sterilized distilled water daily for three months. The remaining groups 2, 3 and 4, were treated daily with 20 mg/kg, 100 mg/kg and 500 mg/kg, respectively of APE suspended in sterilized distilled water for three months. The animals in each group were weighed on day zero (baseline) and weekly thereafter for 3 months.

#### Blood sampling/Clotting time

Blood samples of rats in each group were taken by tail bleeding at the end of the 3 months of treatment into serum separator tubes, centrifuged at 4000 x g for 5 minutes and supernatant collected and stored at −40 °C for biochemical analyses. Other blood samples were collected into separate tubes pre-coated with trisodium citrate for hematological analyses within 24 hrs. Clotting times of other blood samples in Eppendorf tubes were also determined.

#### Urinalysis

Urine samples of the rats in each treatment group produced by involuntary discharges by the groups were collected on clean ceramic tiles at the start of the study (baseline) and after three months of treatment. Analysis of urine for glucose, bilirubin, ketones, specific gravity, pH, proteins, urobilinogen, nitrate, blood and leukocytes were done using urine reagent strips UroColor^TM^ 10 (Standard Diagnostic Inc., Korea).

#### Serum biochemical analysis

Serum alanine transaminase (ALT), alkaline phosphatase (ALP), bilirubin (direct), and creatinine of the samples were determined following the manufacturer’s (ELI Tech) protocol with Biosystems A25 Chemistry Analyzer.

#### Hematological analysis

Red blood cell (RBC) count, white blood cell (WBC) count, hematocrit (HCT), hemoglobin (HGB), mean cell volume (MCV) and platelet count (PLT), mean corpuscular hemoglobin (MCH), mean corpuscular hemoglobin concentration (MCHC), lymphocytes (LYM), red blood cell distribution width (RDWSD), Mean platelet volume (MPV) of blood were determined with Haema-screen 18 (Hospitex Diagnostics, Italy) following established protocol.

#### Pentobarbital-induced sleeping time

The sleep evaluation method was based on the potentiation of pentobarbital-induced sleeping time^15^. Briefly, the four (4) animals in each treatment group were given a single dose of pentobarbital (40 mg/kg) to induce sleep. The time the animals lost the righting reflex till the time they gained the righting reflex was determined. The interval between these two times was taken as the pentobarbital-induced sleeping time.

#### Mean organ wet weights

The animals were weighed and sacrificed at the end of the study by cervical dislocation. The body organs (kidneys, liver, lungs, and heart) of each rat were excised, washed in ice-cold 1.15% KCl solution, dried, and weighed. Organ wet weights were expressed as g% ((g of organ weight/ g of body weight) x100%).

#### Histopathological analysis

At the end of the 3 months treatment time, the animals were euthanized humanely, and the liver, kidneys, lungs, and heart were surgically removed and fixed in 10% formalin. The organs were dehydrated in varying concentrations of ethanol (70-100%) and then cleared in xylene before embedding in paraffin wax. Micro-sections (4 μm) were prepared, mounted on slides, and stained with hematoxylin and eosin (H&E) and examined under a light microscope by a pathologist in a blinded fashion ^16^.

#### Statistical Analysis

Results were presented as mean ± standard error of the mean (SEM) for n of at least 6 observations. Statistical analysis was done using one-way analysis of variance (ANOVA) followed by Dunnett post hoc analysis using Graph Pad Prism 5.0. Statistical significance was considered at p value of less than 0.05.

## RESULTS

### Acute Toxicity of Annickia polycarpa extract

The effect of a single oral dose of 5000 mg/kg APE administered to six animals did not result in any deaths within 24 h, and no physical signs of toxicity as evidenced by normal breathing and locomotion, and absence of signs of tremors, convulsions, salivation, diarrhea, lethargy, excessive lachrymatory activity, and piloerection over the 14 days of study. Hence, the LD50 of APE is higher than 5000 mg/kg and is relatively non-toxic.

### Sub-chronic toxicity studies of *Annickia polycarpa* extract

*Annickia polycarpa extract did not cause changes in body weight over time* Changes in body weight have been linked to adverse effects of chemicals and drugs. We therefore monitored weight changes throughout the 3 months of treatment. There were significant increases in rat mean body weight changes with time over the experimental period for all animal treatment groups. However, there were no significant differences between the APE-treated groups and the control group (Fig. 1).

**Figure 1:**
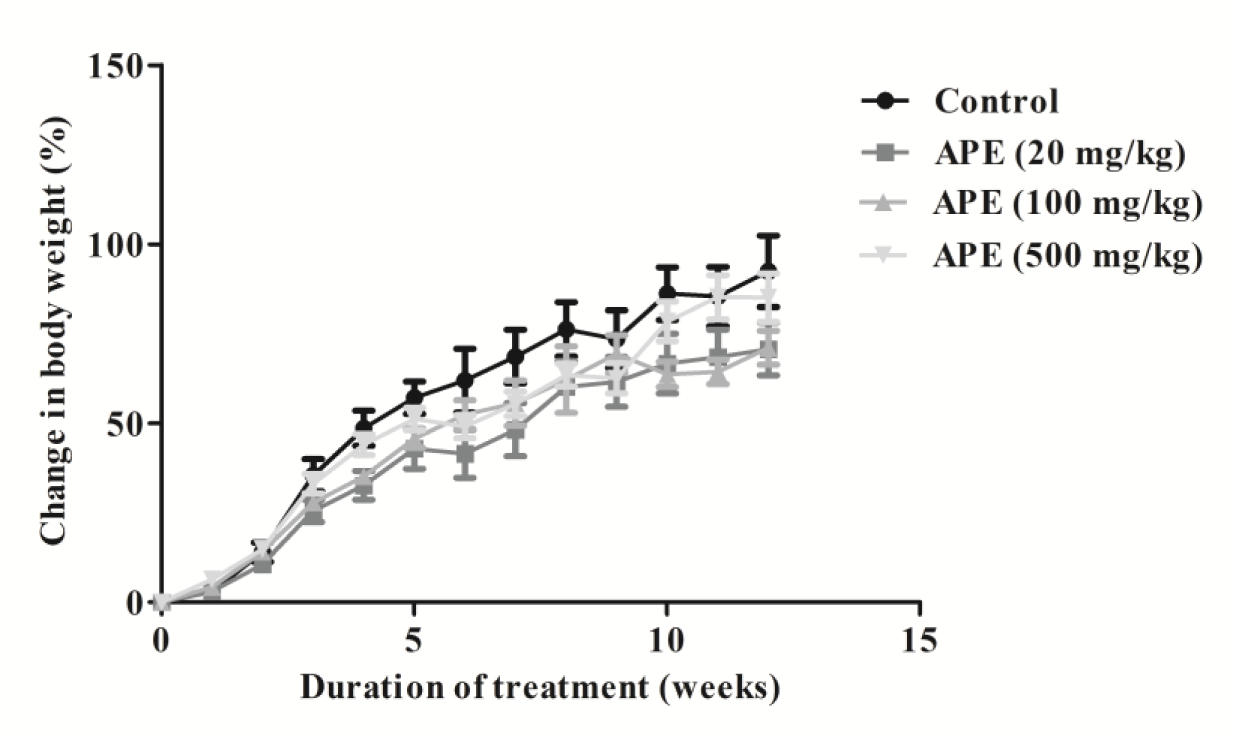
Percentage changes in body weight of Sprague-Dawley rats over duration of treatment with APE. Values are expressed as percentage change in baseline control value of 131.5 ± 1.2 g. Results are mean ± S.E.M of n=6. APE= *A. polycarpa* extract.

In addition, we determined the effect of treatment with APE on organ wet weights expressed as a percentage of body weight. There were no significant differences in the wet weights of organs (heart, kidney, liver, lung, and spleen) between the APE-treated groups and the control group (Table 1). Thus, APE administration to mice does not cause changes in organ wet weight as well as body weights.

**Table 1:** Effect of treatment with APE on wet organ/body weight.

| Organ wet weights/body weight (%) |  |  |  |  |
| --- | --- | --- | --- | --- |
| Parameters | Treatment Groups |  |  |  |
|  | Control | APE (20 mg/kg) | APE (100 mg/kg) | APE (500 mg/kg) |
| Heart | 0.29 ± 0.01 | 0.27 ± 0.01 | 0.29 ± 0.01 | 0.28 ± 0.01 |
| Kidney | 0.49 ± 0.03 | 0.51 ± 0.03 | 0.49 ± 0.01 | 0.49 ± 0.02 |
| Liver | 2.9 ± 0.23 | 2.84 ± 0.12 | 2.94 ± 0.14 | 2.67 ± 0.27 |
| Lung | 0.66 ± 0.06 | 0.61 ± 0.02 | 0.58 ± 0.03 | 0.73 ± 0.11 |
| Spleen | 0.25 ± 0.02 | 0.22 ± 0.02 | 0.23 ± 0.01 | 0.23 ± 0.02 |

552 Values are means ± S.E.M. of n=6. APE= *A. polycarpa* extract

#### Annickia polycarpa extract did not cause changes in hematological indices

Drug use, especially in a prolonged manner, has a tendency to cause changes in blood parameters such as red blood cell count, white blood cell count, and platelet count. All hematological parameters measured after 3 months of treatments indicated that there were no significant differences in values between APE-treated groups at all dose levels and the control group (Table 2). APE therefore does not pose any hematotoxic effect.

**Table 2:** Effect of APE on haematological parameters at termination of treatments.

| Parameters | Treatment Groups |  |  |  |
| --- | --- | --- | --- | --- |
|  | Control | APE (20 mg/kg) | APE (100 mg/kg) | APE (500mg/kg) |
| WBC (10<sup>9</sup>/l) | 12.87 ± 1.90 | 14.57 ± 1.10 | 12.50 ± 0.93 | 14.57 ± 1.09 |
| RBC (10<sup>12</sup>/L) | 8.78 ± 0.32 | 8.80 ± 0.42 | 8.46 ± 0.18 | 8.83 ± 0.30 |
| HGB (g/dl) | 14.70 ± 0.10 | 15.43 ± 0.67 | 15.00 ± 0.46 | 14.90 ± 0.26 |
| HCT (%) | 46.97 ± 0.82 | 49.00 ± 2.52 | 47.23 ± 1.37 | 47.33 ± 1.11 |
| MCV (fl) | 53.60 ± 1.27 | 55.67 ± 0.43 | 55.83 ± 0.47 | 53.63 ± 0.64 |
| MCH (pg) | 16.80 ± 0.55 | 16.57 ± 1.19 | 17.70 ± 0.17 | 16.87 ± 0.29 |
| MCHC (g/dl) | 31.30 ± 0.35 | 31.53 ± 0.29 | 31.77 ± 0.15 | 31.50 ± 0.20 |
| PLT (10<sup>9</sup>/l) | 1193 ± 102 | 922.0 ± 6.0 | 1020 ± 116 | 990.7±87.6 |
| LYM(%) | 67.00 ± 1.35 | 67.76 ± 1.47 | 69.93 ± 2.60 | 68.33 ± 4.33 |
| LYM# | 8.60 ± 1.21 | 11.93 ± 0.99 | 8.70 ± 0.66 | 9.83 ± 0.20 |
| RDWSD (fl) | 30.60 ± 0.20 | 28.70 ± 1.03 | 24.30 ± 4.62 | 28.40 ± 0.29 |
| MPV (fl) | 7.47 ± 0.07 | 8.20 ± 0.10 | 7.85 ± 0.15 | 7.87 ± 0.23 |

(RBC= Red blood cell count, WBC= white blood cell count, HCT= hematocrit, HGB= hemoglobin, MCV= mean cell volume, PLT= platelet count mean, MCH= corpuscular hemoglobin, MCHC= mean corpuscular hemoglobin concentration, LYM= lymphocytes, RDWSD= Red blood cell distribution width, MPV=Mean platelet volume). Values are mean ± SEM of n=3. APE= *A. polycarpa* extract.

#### Annickia polycarpa extract did not cause pathological changes in urine

The dipstick urinalysis test is used to detect pathological changes in urine. Dipstick urinalysis of control and test animals at the termination of treatments showed no differences in urine parameters such as specific gravity, proteins, and pH (Table 3). Therefore, the prolonged use of APE does not cause pathological changes in urine.

**Table 3:**
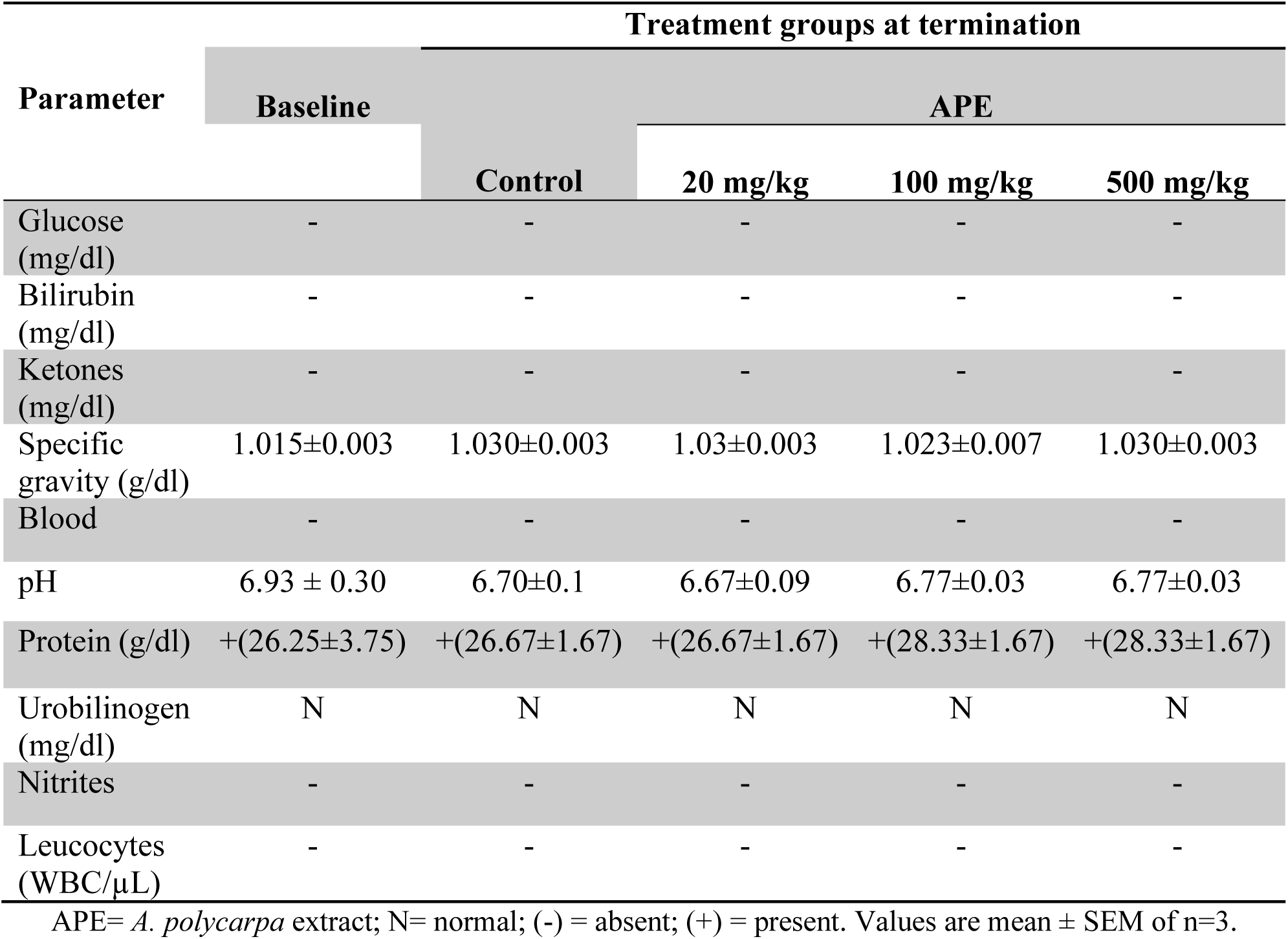
Effect of treatment with APE on urine parameters at baseline and termination of treatments.

#### Effect of Annickia polycarpa extract on liver tissue morphology and function

Having shown that APE treatment does not affect the liver wet weight, we thought that APE may not have any effect on the function of the liver. We therefore determined the effect of APE treatment on liver function. Serum levels of alkaline phosphatase (ALP) and alanine transaminase (ALT) are used as indices to assess liver function. We showed that APE treatment did not significantly affect the serum levels of ALP (Fig. 2A) and ALT (Fig. 2B) at all the doses tested.

**Figure 2:**
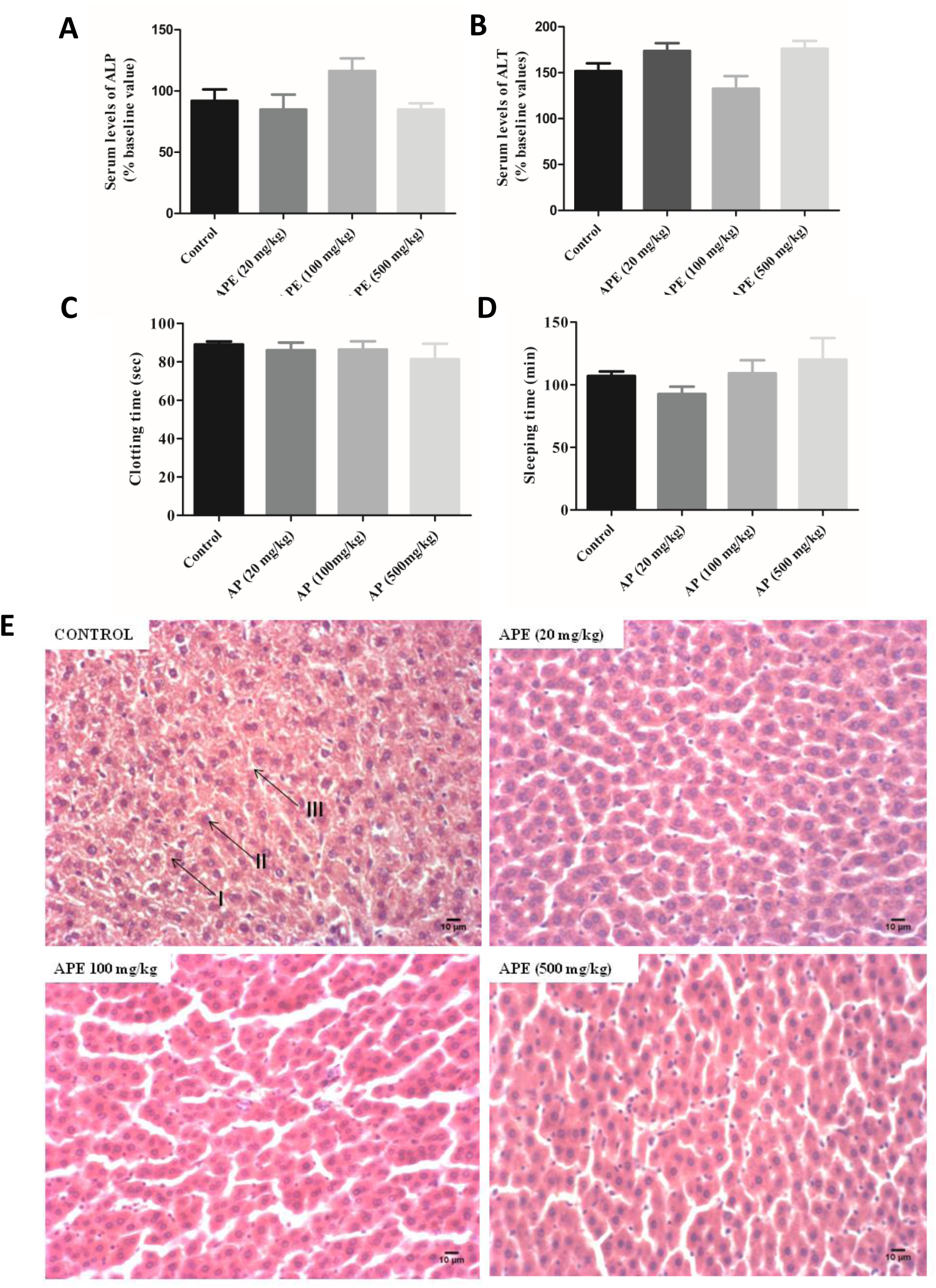
Effect of *Annickia polycarpa* extract on liver tissue morphology and function. (A) Serum Alkaline phosphatase (ALP). Values are percentages of baseline control of ALP (440.4 ± 29.2 U/L), (B) Serum Alanine transaminase (ALT). Values are percentages of baseline control of ALT (53.52 ± 4.57 U/L), (C) pentobarbital-induced sleeping time, and (D) blood clotting time determined at the end of the 3 months treatment with APE. (E) Representative histological images of rat liver at termination of treatments. The control group and the APE-treated groups all show normal histomorphology. normal hepatocyte (I), nuclei (II) and interstitial spaces (III). APE= *A. polycarpa* extract. Magnification: x200. Results mean ± SEM of n = 6. * Value significantly (p<0.05) different from the control group. APE= *A. polycarpa* extract.

Most blood clotting factors are synthesized by the liver. As an indirect measure of another function of the liver, we determined the time it will take for blood from both control groups and APE-treated groups to clot. The blood clotting times of animals treated with APE were not significantly different from control animals at the termination of treatments (Fig. 2D). Also, damage to the liver would affect the levels of cytochrome P450 and consequently sleep time. Again, there were no significant differences in the sleep time between APE-treated animals and controls at the termination of treatments (Fig. 2D).

To corroborate our functional studies of the liver, we examined H&E-stained sections of the liver at the end of treatment. The micrographs of the liver showed normal tissue histomorphology and controls were comparable to the APE-treated groups (Fig 2E). Altogether, APE treatment in a sub-chronic use does not pose any hepatotoxic effect at the dose levels used.

#### Effect of Annickia polycarpa extract on kidney tissue morphology and function

With no pathological changes as revealed by the urinalysis test, we reasoned that APE may not be having any effect on the normal functioning of the kidney. Thus, we determined serum levels of creatinine and direct bilirubin as tests for kidney function. There were no significant differences in serum levels of creatinine and direct bilirubin between controls and the APE-treated animals at all dose levels (Fig. 3A and 3B).

**Figure 3:**
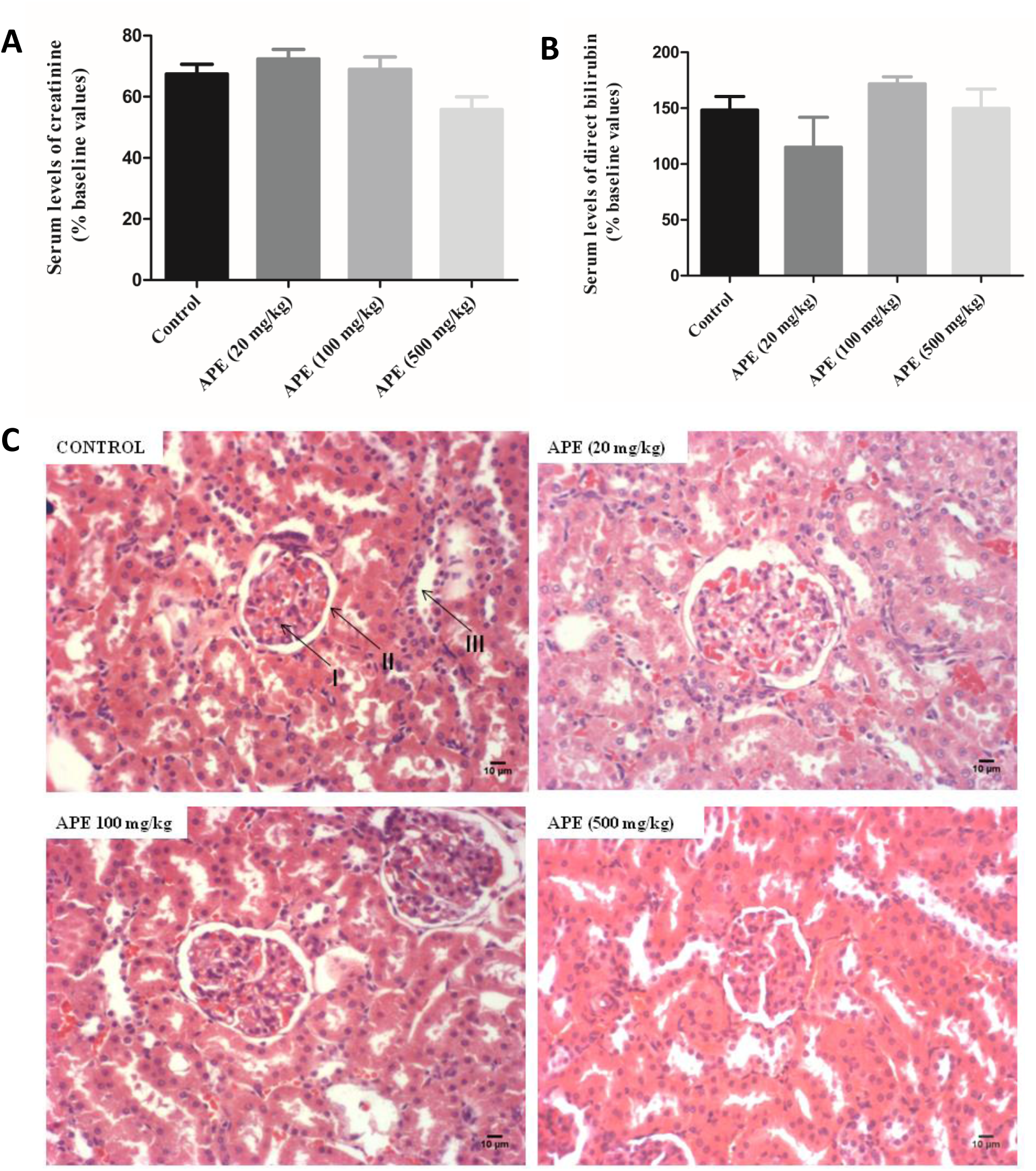
Effect of Annickia polycarpa extract on kidney tissue morphology and function. (A) Effect of APE on serum levels of creatinine. Values are percentages of baseline control of (0.97 ± 0.05) mg/dl, and (B). Serum levels of direct bilirubin at the termination of treatment. Values are percentages of baseline control (0.23 ± 0.02 mg/dl). (C) Representative histological appearance of rat kidney at termination of treatments showing no difference in appearance of (I) glomerulus, (II) bowman’s capsule and (III) renal tubules in both control animals and APE-treated animals. APE= *A. polycarpa* extract. Magnification: x100. Results are mean ± SEM. * Value significantly (p<0.05) different from the control group. APE= *A. polycarpa* extract.

We additionally analyzed the kidneys of APE-treated animals and controls after H&E staining. Both control and APE-treated animals showed normal kidney morphology seen as normal glomerular areas and tubular areas without any sign of edema or leukocyte infiltration (Fig 3C). Thus, APE is not nephrotoxic.

#### Effect of Annickia polycarpa extract on the lung and heart tissue morphology

Generally, the lungs of both control animals and those of APE-treated groups were normal and showed no signs of inflammation or tissue damage. However, there were mild morphological changes in the lung of the APE-treated group in a dose dependent manner characterized by interstitial fibrosis and thickening of alveolar septa, with alveolar space closing (Fig 4A).

**Figure 4:**
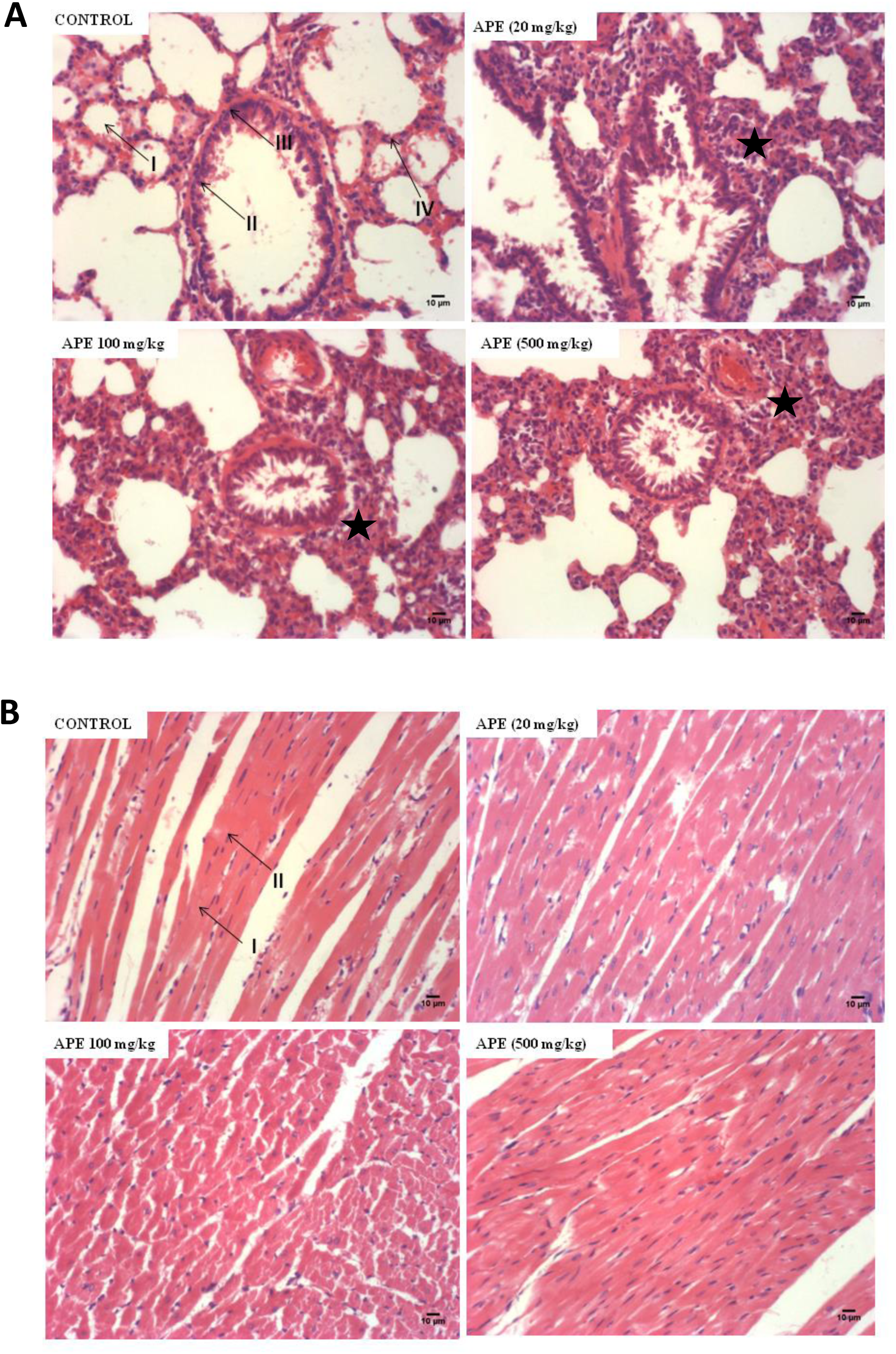
Effect of Annickia polycarpa extract on lung and heart tissue morphology and function. (A) Representative histological appearance of rat lung at termination of treatment. Control animals show normal alveolar areas (I) with Clara cells (II) lining the bronchiolar epithelium (III), and normal interstitial space (IV). Animals treated with APE show a dose-dependent lung damage characterized by interstitial fibrosis and thickening of alveolar septa (black star) and alveolar space closing. (B) Representative histological appearance of rat heart muscle at termination of treatments showing no differences in morphology of (I) cardiac muscle fibres and (II) nuclei of both control and APE-treated animals. APE= *A. polycarpa* extract. Magnification: x200 APE= *A. polycarpa* extract. Magnification: x100

In addition, the micrographs of the heart of rats observed at termination of treatment with APE showed no observable morphological differences compared with the control animals (Fig 4B). Together, while APE showed no toxic effects on the heart, there was selective toxicity on the lungs.

## DISCUSSION

This study was conducted to determine the acute and sub-chronic toxicity of aqueous stem bark extract of *Annickia polycarpa* (APE) in Sprague-Dawley rats. Our laboratory has previously determined the antidiabetic, antioxidant, anti-inflammatory and anti-colitis effects of the APE ^9,14^. Also, APE and its isolated constituents have been found to relieve pain and possessed anti-inflammatory and analgesic effects ^13^. We showed that APE is generally non-toxic except for mild toxic effect on the lung when used over a prolonged period.

The acute toxicity studies showed a median lethal dose (LD50) of APE in rats greater than 5000 mg/kg, with no physical signs of toxicity, an indication that it is relatively safe. This agrees with similar work that reported no mortality at dosage levels of 2000 mg/kg, 4000 mg/kg and 5000 mg/kg during acute toxicity testing ^7,13^. Adverse changes in body weight have been used as an indicator of toxicity of drugs and chemicals ^17^. The normal growth of APE-treated animals in this study, therefore, suggests the absence of toxicity (Fig. 1). Adverse changes in mean organ weight are also a reflection of organ-specific toxicity. Treatment with APE did not affect mean organ/body weights (Table 1) suggesting organs were not edematous as a result of defects in osmoregulation or hypertrophic as a result of lack of control of cell growth and division ^18,19^.

Blood is an essential circulatory tissue that consists of red blood cell, white blood cells and platelets suspended in plasma and functions to maintain homeostasis ^20,21^. Red blood cells serve as carriers of hemoglobin which transports oxygen to tissues ^22^. A reduced red blood cell count suggests that oxygen and carbon dioxide transport will be impaired ^22^. White blood cells help to fight infections, and as a result, reduced white blood cell count makes one susceptible to infections ^22,23^. Blood platelets are associated with blood clotting, and for that matter, low platelet concentration will lead to long clotting time leading to blood loss ^24–26^. The lack of effect of APE in platelet count is reflected in its lack of effect on blood clotting time. Some haemotoxicants, for example, paracetamol and some phytochemicals are known to cause a reduction in red blood cell counts and also alter hemoglobin concentration with subsequent anemia in experimental animals ^27^. Oral administration of APE for three months did not result in a significant difference between hematological parameters of the control and treatment groups. This presupposes that the extract has no direct effect on the blood cells or bone marrow or indirect effect on erythropoiesis.

The liver is a key organ in the body involved in metabolism, storage, and excretion of exogenous compounds as well as the metabolism of endogenous compounds ^28,29^. It is the powerhouse of the body ^30^. Toxic insults to the liver, causes damaged hepatocytes to release enzymes such as alkaline phosphatase, alanine aminotransferase and aspartate aminotransferase into the blood, thus increasing the serum levels of these enzymes ^31–34^. Toxic insults to the liver also affect its biosynthetic ability, thus leading to impaired protein synthesis. The liver is the site of synthesis of blood clotting factors ^35^, which are protein in nature; hence damage to the liver will prolong blood clotting time (prothrombin time) ^36^. This study showed that APE did not cause the elevation of these serum enzyme levels and did not affect the blood clotting time, an indication of the absence of hepatocellular damage. These observations are corroborated by the absence of adverse changes in the morphology of liver cells and, additionally, the lack of effect on pentobarbital induced sleeping time in APE-treated animals. Damage to the liver cells may result in the reduction of cytochrome P450 levels, thus leading to an increase in pentobarbital induced sleeping time as a result of reduced metabolism of pentobarbital ^15,37^.

The kidneys produce erythropoietin which is involved in erythropoiesis, and therefore damage to the kidney impairs erythropoiesis and blood cell formation ^38^. Damage to the glomerulus causes an increase in serum creatinine and urea levels as well as the presence of albumin in the urine ^39^.

The renal tubules are responsible for concentrating urine, regulation of ion excretion and the production of acidic urine ^39^. There were no significant differences in urine protein and pH between baseline values and termination values for control and extract-treated groups. The extract did not cause any changes in the serum and urine parameters and did not affect blood cell formation. This seems to suggest that the extract may not be nephrotoxic. These results are corroborated by the morphology of the kidney, which shows no adverse effects on the Bowman’s capsule, glomerulus, and renal tubules (Fig. 3C).

The heart is a muscular organ which functions by pumping blood to all parts of the body, supplying tissues with oxygen and nutrients ^40^. The lung is the site for gaseous exchange in the body. APE did not cause any morphological changes in the heart muscle, indicating that it may not be cardiotoxic (Fig. 4B). However, in the case of the lung, APE appeared to cause alveolar cell damage, without causing bronchiolar cell damage, which appears to be dose-dependent (Fig. 4A) and characterized by the thickening of the alveolar septa and interstitial fibrosis. This may affect the diffusion of oxygen across the alveoli with its attendant effect on the degree of oxygen transport and thus, cellular metabolism. The extract thus appears to be mildly pneumotoxic so care should be taken in its use.

## CONCLUSION

*A. polycarpa* extract has no hepatotoxic, nephrotoxic or cardiotoxic effects, but care must be taken in its use because of its mild selective lung toxicity when used continuously for a prolonged period of time.

## DATA AVAILABILITY

The data used to support the findings are included within the article.

## ACKNOWLEDGEMENTS

This work was part of MPhil research carried out in the Department of Chemical Pathology, University of Ghana Medical School by Nathaniel L. Lartey. Nathaniel’s master’s program was partially supported by a scholarship by the College of Health Sciences, University of Ghana. The authors are grateful to Mr Donkor for his immense help in taking micrographs of the tissue cryosections. The authors are also grateful to the staff of the Animal Experimentation Unit of the CPMR and the Electron Microscopy Department of the Noguchi Memorial Institute for Medical Research.

